# Identification of novel common variants associated with chronic pain using conditional false discovery rate analysis with major depressive disorder and assessment of pleiotropic effects of *LRFN5*

**DOI:** 10.1101/495846

**Authors:** Keira J.A. Johnston, Mark J. Adams, Barbara I. Nicholl, Joey Ward, Rona J Strawbridge, Andrew McIntosh, Daniel J. Smith, Mark E.S. Bailey

## Abstract

Chronic pain is highly prevalent worldwide, with a significant socioeconomic burden, and also contributes to excess mortality. Chronic pain is a complex trait that is moderately heritable and genetically, as well as phenotypically, correlated with major depressive disorder (MDD). Use of the Conditional False Discovery Rate (cFDR) approach, which leverages pleiotropy identified from existing GWAS outputs, has been successful in discovering novel associated variants in related phenotypes. Here, genome-wide association study outputs for both von Korff chronic pain grade as a quasi-quantitative trait and for MDD were used to identify variants meeting a cFDR threshold for each outcome phenotype separately, as well as a conjunctional cFDR (ccFDR) threshold for both phenotypes together. Using a moderately conservative threshold, we identified a total of 11 novel single nucleotide polymorphisms (SNPs), six of which were associated with chronic pain grade and nine of which were associated with MDD. Four SNPs on chromosome 14 were associated with both chronic pain grade and MDD. SNPs associated only with chronic pain grade were located within *SLC16A7* on chromosome 12. SNPs associated only with MDD were located either in a gene-dense region on chromosome 1 harbouring *LINC01360, LRRIQ3, FPGT* and *FPGT-TNNI3K*, or within/close to *LRFN5* on chromosome 14. The SNPs associated with both outcomes were also located within *LRFN5*. Several of the SNPs on chromosomes 1 and 14 were identified as being associated with expression levels of nearby genes in the brain and central nervous system. Overall, using the cFDR approach, we identified several novel genetic loci associated with chronic pain and we describe likely pleiotropic effects of a recently identified MDD locus on chronic pain.

**Author Summary:** Genetic variants explaining variation in complex traits can often be associated with more than one trait at once (‘pleiotropy’). Taking account of this pleiotropy in genetic studies can increase power to find sites in the genome harbouring trait-associated variants. In this study we used the suspected underlying pleiotropy between chronic pain and major depressive disorder to discover novel variants associated with chronic pain, and to investigate genetic variation that may be shared between the two disorders.

## Introduction

Chronic pain is defined as pain lasting longer than 12 weeks. It affects around 30% of the global adult population (Elzahaf, Tashani, Unsworth, & Johnson, 2012), imposes significant socioeconomic burden, and contributes to excess mortality (Hocking, Morris, Dominiczak, Porteous, & Smith, 2012; Vos et al., 2015). Chronic pain is often associated with both specific and non-specific medical conditions (such as cancers and HIV/AIDS, fibromyalgia and musculoskeletal conditions), and with a range of injuries (Greene, 2010; Merskey & Bogduk, 1994). It is also commonly co-morbid with mood disorders, such as major depressive disorder (MDD; Flint & Kendler, 2014; McIntosh et al., 2016). MDD is a common and serious mood disorder involving psychological symptoms such as persistent low mood and anhedonia, and physical symptoms such as changes in appetite and sleep disturbance (American Psychiatric Association 2013). In common with chronic pain, MDD also imposes a significant socioeconomic burden worldwide and is now the leading cause of disability globally (Vos et al., 2015). The causal direction of factors underlying the association between chronic pain and mood disorders is unclear. Chronic pain is a complex trait with moderate heritability (Hocking et al., 2012) and chronic pain and MDD are genetically correlated (McIntosh et al., 2016), suggesting that pleiotropic variants exist that predispose to both conditions. To date, genome-wide association studies (GWAS) have not identified any genome-wide significant variants associated with chronic pain at any site (Mogil, 2012; Zorina-Lichtenwalter, Meloto, Khoury, & Diatchenko, 2016).

An alternative strategy for identifying genetic variation contributing to complex traits is to make use of existing GWAS summary statistic outputs within more powerful discovery approaches, such as the conditional false discovery rate (cFDR) approach (Andreassen, Thompson, et al., 2013). The cFDR approach makes better use of the genetic overlap between traits or conditions to discover loci based on their pleiotropic effects in each phenotype separately, as the strength of evidence for involvement of a locus in a primary trait may be enhanced by consideration of the strength of evidence for the same locus in an associated secondary trait. In the case of chronic pain, the association with mood disorders and mood-related traits is of substantial interest, as improved understanding of the biological underpinnings of this overlap may stimulate the development of novel treatment strategies.

cFDR analyses have been used to find novel variants associated with schizophrenia, type 2 diabetes, Alzheimer disease, bipolar disorder and systolic blood pressure (Andreassen, Djurovic, et al., 2013; Andreassen et al., 2014; Andreassen, Thompson, et al., 2013; Wang et al., 2016). This therefore represents a promising and potentially more cost-effective method for identifying new SNPs associated with complex traits by maximising the utility of existing GWAS outputs. We aimed to find chronic pain-associated SNPs using cFDR analysis, with MDD as the secondary trait. In addition, we sought to discover loci with pleiotropic effects on these two phenotypes. Our findings revealed six novel chronic pain-associated SNPs and a locus within the *LRFN5* gene where the nature of the pleiotropic effects is consistent with major depression risk being mediated by genetic effects on risks of chronic pain. These findings contribute to our understanding of the biological underpinnings of chronic pain and a component of its shared biology with major depressive disorder.

## Methods

### Phenotype definition and source data

For chronic pain in the cFDR analysis, summary statistics from a GWAS carried out collaboratively with Pfizer-23andMe Inc (McIntosh et al., 2016) were used. In that study, chronic pain severity was modelled as a quasi-quantitative trait, defined by von Korff and colleagues as ‘chronic pain grade’ (CPG; Von Korff, Ormel, Keefe, & Dworkin, 1992), on a scale from 0 (no chronic pain reported) to 4 (high intensity, high disability, severely limiting chronic pain). The GWAS was carried out using data from 23, 301 unrelated individuals of white European descent (10, 543 individuals with CPG levels 1-4 combined, 12, 758 individuals with CPG level 0; Supplementary Table 1). For Major Depressive Disorder (MDD), summary statistics from a recent case-control GWAS meta-analysis (Wray et al., 2018) were provided by the Psychiatric Genomics Consortium (PGC) after removal of data from 23andMe and UK Biobank participants, yielding a dataset originating from an analysis using 43,028 cases and 87, 522 controls. Thus, none of the participants contributing to the CPG summary statistics overlapped with those contributing to the MDD summary statistics. Phenotype definitions, study population demographics and meta-analysis procedures for the MDD GWAS have been described previously (Wray et al., 2018).

### Data Preparation & LD Pruning

A dataset of SNPs for which a *p*-value for association, chromosome position data and rsID were available in both MDD and CPG datasets was compiled. This was then LD pruned as follows. Firstly, PLINK-format genotype data, for each SNP in the newly compiled CPG-MDD summary-statistic dataset, was extracted from the UK Biobank genotype data (approved application 6553). Pruning was carried out using command line PLINK (version 1.9) --indep-pairwise function. From each SNP-pair with r^2^ > 0.2 within a 50-SNP window of the lead SNP, one was removed according to PLINK’s greedy pruning algorithm, the window slid along 5 SNPs, and the pruning repeated. This resulted in an LD-pruned dataset of 774,292 SNPs with association data available for both MDD and CPG.

### Conditional False Discovery Rate (cFDR) analyses

cFDR is an extension of local-area false-discovery rate (FDR), which is in turn a re-thinking of Benjamini & Hochberg’s ‘tail-end’ false discovery rate procedure (Benjamini & Hochberg, 1995; Efron, 2007; Liley & Wallace, 2015; Storey, 2002). The tail-end FDR procedure is concerned with controlling the FDR at a pre-defined level and deciding the maximum test statistic value from a list of ordered test statistic values that allows for this (Benjamini & Hochberg, 1995), which becomes the cut-off value for deciding significance. Local FDR reframes the FDR as a Bayesian posterior probability that the SNP in question is not associated with the disease or trait, given its association test statistic (usually its *p* value) (Efron, 2007; Storey, 2002).

cFDR analysis of GWAS summary statistics simply extends local FDR, with the result being the posterior probability that the SNP in question is not associated with the primary phenotype, given its association test statistic value for both the primary phenotype and for a secondary, related phenotype. This allows for exploitation of any underlying pleiotropic genetic architecture, believed to be ubiquitous in human complex disease (Gratten & Visscher, 2016; Sivakumaran et al., 2011), along with an increase in power compared to traditional Bonferroni correction of GWAS summary statistic output.

cFDR and conjunctional cFDR (ccFDR) values for each SNP for CPG given MDD and vice versa were calculated (Formula 1) as previously detailed (Liley & Wallace, 2015) using the statistical software R (version 3.3.3).

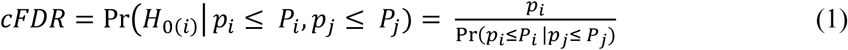

The cFDR value for a SNP obtained from the above formula is the probability that the SNP is not associated with the primary phenotype, given its strength of association with the secondary phenotype. The ccFDR was obtained via taking the higher of the two cFDR values obtained for each SNP. The significance threshold chosen was cFDR (and ccFDR) ≤0.01.

### Genomic Context Analyses

The genomic context for SNPs with significant cFDR values in either analysis was examined. The R package ‘rsnps’ (Chamberlain et al 2016) was used to extract data from records in NCBI dbSNP (https://www.ncbi.nlm.nih.gov/snp). Gene context for each SNP was examined in the UCSC Genome Browser (build GRCh38/hg38) (Kent, Sugnet, Furey, & Roskin, 2002), using a window of 0.5Mbp around each SNP and data from the GENCODE v24 track, validated or reviewed by either Refseq or SwissProt staff. Thepresence of cis-eQTLs close to the significant SNPs was investigated using the IGV eQTL Browser (Aguet et al., 2017) web interface.

### Further Exploration of Pleiotropy in *LRFN5*

Regression models were run to examine associations between genotype at a SNP identified as pleiotropic in the ccFDR analysis and relevant phenotypes in data from the UK Biobank (chronic pain, MDD, age attended assessment centre and sex; approved applications 10302, 6553). Genotype at rs11846556 (the SNP with the lowest ccFDR) in UK Biobank individuals was coded as number of effect-associated alleles (0, 1, 2 copies of the A allele).

Chronic pain was assessed via the question ‘pain types experienced in the past month’, with possible answers as follows: each of 7 different body sites (multiple sites allowed), or ‘all over the body’, or ‘do not know’, or ‘prefer not to answer’. In a separate question, the duration of any pain at the reportable sites/all over the body was assessed. A dichotomous chronic pain variable was defined as the presence of chronic pain in at least 1 site (or all over the body) for longer than 3 months. Those who did not report pain of that duration were assigned to a control group.

In addition to this dichotomous ‘presence of chronic pain’ variable, a chronic pain variable capturing information about the number of sites of chronic pain was also created (chronic pain category; treated as an ordinal variable in regression models), again employing the 3 month minimum duration criterion: 0 = no chronic pain, 1 = single site, 2 = 2-3 sites, 3 = 4-7 sites, 4 = all over the body (Nicholl et al 2014). Individuals giving ‘prefer not to answer’ and ‘do not know’ answers were excluded from all analyses.

MDD cases were ascertained according to Davis et al 2018 from a subset of UK Biobank participants who completed a detailed the mental health questionnaire assessment (N = 157, 366). Controls consisted of those with no mood disorder; those with bipolar disorder were excluded from analyses. An ordinal MDD phenotype (MDD severity) was also obtained by classifying MDD by number of episodes: 0 = no mood disorder, 1 = single episode MDD, 2 = recurrent MDD.

A UK Biobank dataset where all individuals had complete information on age, sex, rs11846556 genotype and chronic pain variables was assembled (N = 469, 253). Logistic regression of chronic pain (case-control) on rs1186556 genotype (adjusted for age and sex) and ordinal regression of chronic pain category on rs1186556 genotype (adjusted for age and sex) were then carried out to assess the relationship between SNP genotype and chronic pain.

A second dataset where all individuals had complete information on age, sex, rs11846556 genotype and MDD variables was assembled (N = 122,286). Logistic regression of MDD (case-control) on rs11846556 genotype (adjusted for age and sex), and ordinal regression of MDD severity on rs11846556 genotype (adjusted for age and sex) were then performed to assess the relationship between SNP genotype and MDD.

A final dataset, where all individuals had complete information on age, sex, genotype, chronic pain and MDD variables was assembled (N = 121,246). Logistic regressions of MDD (case-control) on chronic pain (case-control) and genotype (adjusted for age and sex), and of chronic pain (case-control) on MDD (case-control) and genotype (adjusted for age and sex) were then carried out, to assess whether any relationship between MDD and SNP genotype was attenuated by chronic pain, and whether any relationship between SNP genotype and chronic pain was attenuated by MDD.

P values for ordinal regression estimates were obtained via comparing the t value obtained to that of a normal t-distribution. Standard errors were used to calculate 95% CI values for ordinal regression output. All models were adjusted for age and sex, both of which were highly significant in all cases. Some models were additionally adjusted for the phenotype (pain or MDD) not being tested as the outcome variable.

## Results

### cFDR and ccFDR analyses

The conditional false discovery approach was used to identify variants associated with chronic pain grade, MDD, or both together in a pleiotropic fashion. Eleven SNPs in total were found at cFDR ≤ 0.01 (Table 1), six of which, located on chromosomes 12 and 14, were associated with CPG and nine of which, located on chromosomes 1 and 14, were associated with MDD. Four of these 11 SNPs, all located within a 131 kbp region on chromosome 14, were found to be pleiotropic (ccFDR ≤0.01).

**Table 1:** Loci identified from cFDR analysis. Trait = trait with which SNP is associated at cFDR ≤0.01. Position = position given as chromosome:base pair location. cFDR (CPG) = cFDR for CPG conditioning on MDD; Beta (CPG)/p (CPG) = effect size and p value from the CPG GWAS; cFDR (MDD) = cFDR for MDD conditioning on CPG; OR (MDD)/p (MDD) = effect size (odds ratio for the effect allele) and p value from the MDD GWAS. Alleles are given as effect allele/other; effect allele is defined as the allele for which association with the trait was tested in the original (CPG or MDD) GWAS. Significant p-values and cFDR/ccFDR values are shown in bold.

| Trait | rsID | Position | Alleles<br>(CPG) | Beta (CPG) | P (CPG) | cFDR (CPG) | Alleles<br>(MDD) | OR (MDD) | P (MDD) | cFDR (MDD) | ccFDR |
| --- | --- | --- | --- | --- | --- | --- | --- | --- | --- | --- | --- |
| MDD | rs35641559 | 1:73760104 | C/T | 0.02 | 0.03 | 0.1 | T/C | 1.04 | 2.08E-06 | 8.83E-03 | 0.11 |
| CPG | rs147573737 | 12:60231575 | C/T | -0.2 | 2.09E-07 | 2.21E-03 | T/C | 1.09 | 0.023 | 0.023 | 0.023 |
| CPG | rs149981001 | 12:60264802 | C/T | 0.2 | 6.24E-08 | 1.06E-03 | T/C | 0.92 | 0.0181 | 0.018 | 0.018 |
| MDD | rs10872954 | 14:41948768 | A/G | -0.026 | 6.45E-03 | 0.053 | A/G | 1.04 | 7.68E-06 | 7.02E-03 | 0.053 |
| MDD | rs10138559 | 14:41975989 | C/T | -0.02 | 0.03 | 0.1 | T/C | 0.96 | 1.04E-06 | 5.25E-03 | 0.1 |
| Both | rs11157241 | 14:42051771 | C/T | 0.035 | 2.91E-04 | 5.83E-04 | T/C | 1.06 | 4.44E-09 | 1.28E-06 | 5.83E-04 |
| Both | rs8015100 | 14:42095232 | A/T | -0.033 | 6.67E-04 | 6.67E-04 | A/T | 1.06 | 1.50E-09 | 9.27E-07 | 6.67E-04 |
| Both | rs10131184 | 14:42166111 | A/G | 0.035 | 2.82E-04 | 8.46E-04 | A/G | 0.95 | 2.53E-08 | 7.10E-06 | 8.46E-04 |
| Both | rs11846556 | 14:42183025 | A/G | -0.037 | 1.11E-04 | 5.57E-04 | A/G | 1.05 | 2.98E-07 | 3.76E-05 | 5.57E-04 |
| MDD | rs1584317 | 14:42213816 | C/G | 0.028 | 4.48E-03 | 0.029 | C/G | 0.96 | 4.59E-06 | 3.57E-03 | 0.03 |
| MDD | rs4904790 | 14:42242623 | C/T | -0.03 | 1.44E-03 | 0.02 | T/C | 0.96 | 1.37E-05 | 3.58E-03 | 0.02 |

### Genomic Context

The SNPs associated with CPG only were located close to *SLC16A7* on chromosome 12 (Supplementary Table 2a, 2b), while the SNPs associated with MDD only were located within a gene-dense region on chromosome 1 containing *LINC01360, LRRIQ3, FPGT, FPGT-TNNI3K,* amongst other genes (Supplementary Table 2b) or close to (upstream of) or within *LRFN5* on chromosome 14 (Supplementary Table 2a, 2b). The SNPs found to be pleiotropic were also located within *LRFN5* on chromosome 14 (Supplementary Table 2a, 2b). The pleiotropic *LRFN5* SNPs were all located just upstream of the 5’-most promoter, or within a large intron close to the 5’-end of the gene. The SNPs associated with MDD only were also located within this intron, or were located a little further upstream of the gene. The CPG-only SNPs were all located downstream of *SLC16A7*. One of the pleiotropic SNPs on chromosome 14, rs11157241, is located approx. 30 kbp upstream of *LRFN5*.

GTeX eQTL lookups revealed that some of these phenotype-associated SNPs were also associated with expression levels of nearby genes. A SNP associated only with MDD, on chromosome 1 (rs35641559), was found to be significantly associated with expression of a long non-coding RNA gene *LINC01360* in the testis (FDR < 0.05, Supplementary Table 2c). The MDD-associated and pleiotropic SNPs on chromosome 14 are all significantly associated with expression of *LRFN5* in a range of tissues, including brain, heart, adipose tissue and spleen (Supplementary Table 2c). The CPG-associated SNPs on chromosome 12 were not significantly associated with expression of any gene in the eQTL database (Supplementary Table 2c).

### Association between rs11846556 genotype and MDD and chronic pain in UK Biobank

To investigate the nature of the pleiotropy revealed for variants within *LRFN5,* regression models were used to assess the strength of association between rs11846556 genotype and each of the two phenotypes of interest, chronic pain and MDD. Genotype at this SNP was associated with significantly increased risk for chronic pain (Table 2a; FDR-adjusted *p* = 0.0008), with each additional copy of the A allele increasing the odds of having chronic pain by 2% (OR = 1.02). rs11846556 genotype was also associated with chronic pain category (Table 2b), with each additional copy of the A allele associated with an OR of 1.02 per chronic pain category (FDR-adjusted *p* = 4.4×10^-5^). Genotype at rs11846556 was not significantly associated with MDD (Table 2a) or with MDD severity (Table 2b). rs11846556 genotype accounted for 0.29% of the variance in chronic pain presence and 0.25% of the variance in chronic pain category (estimated using McFadden’s R^2^, adjusting for age & sex). rs11846556 genotype was no longer associated with presence of chronic pain after adjustment for MDD status (FDR-adj *p* = 0.096), and with a larger effect size (per allele betas were 0.017 vs 0.015 in the MDD-adjusted and non-adjusted models, respectively).

**Table 2:** Analysis of associations between rs11846556 genotype and chronic pain/MDD.] a) Legend: a) results of logistic regression models with dichotomous chronic pain (CP(dichot.)) and major depressive disorder (MDD) variables as outcomes and SNP genotype as predictor. SE = standard error of the beta coefficient value, OR = odds ratio. Ns = non-significant (p > 0.05).b) results of ordinal regression models with chronic pain category and MDD severity variables as outcomes and SNP genotype as predictor. SE = standard error of the beta coefficient, ns = non-significant (p > 0.05).

|  | **Beta** | **SE** | **OR** | ***p*** | **FDR-**<br>**adj. *p*** | **Additional**<br>**adjuster** |
| --- | --- | --- | --- | --- | --- | --- |
| **CP**<br>**(dichot.)** | 0.015 | 0.004 | 1.02 | 2.63E-04 | 7.89E-04 |  |
| **CP**<br>**(dichot.)** | 0.017 | 0.008 | 1.02 | 0.048 | 0.096 | **MDD**<br>**(dichot)** |
| **MDD** | 0.014 | 0.009 | 1.01 | ns |  |  |
| **MDD** | 0.011 | 0.010 | 1.01 | ns |  | **CP**<br>**(dichot)** |

**b)**
|  | **Beta** | **SE** | ***p*** | **FDR-**<br>**adj. *p*** |
| --- | --- | --- | --- | --- |
| **CPC** | 0.018 | 0.004 | 7.26E-06 | 4.36E-05 |
| **MDD severity** | 0.014 | 0.009 | ns |  |

## Discussion

### Genetics of Chronic Pain & Related Disorders

The genetics of pain have hitherto mostly been investigated using candidate gene and animal-model approaches. No genome-wide significant loci have been reported from GWAS of presence of chronic pain at any site, or of chronic pain severity, although GWAS have been carried out for disorders with chronic pain as a symptom, and loci have been reported for particular subtypes of chronic or regular pain e.g. migraine, back pain and other musculoskeletal disorders (Suri et al., 2018; Zorina-Lichtenwalter et al., 2016). Chronic pain reported at particular sites may be triggered by environmental insults (such as surgery or physical trauma) or by long-term inflammatory processes, but the mechanisms that allow pain to persist are thought to be largely neurogenic in origin, with aberrant feedback between the spinal cord and brain strongly implicated (Apkarian, Hashmi, & Baliki, 2011; Q. Xu & Yaksh, 2011). Thus, exploration of the molecular mechanisms underlying chronic pain is likely to benefit from studies focussing on phenotypes relating to severity and other measures of overall susceptibility to chronic pain.

In this study, the power to detect CPG-associated SNPs was increased by leveraging the additional information coming from the correlation between predisposing loci for CPG and for MDD. Overall, we found six CPG-associated and nine MDD-associated SNPs. In addition, pleiotropic effects of four *LRFN5* SNPs were observed.

### CPG-Associated SNPs

The SNPs associated with CPG only were located close to *SLC16A7,* which encodes monocarboxylate transporter 2 (MCT2). In the central nervous system, MCT2 is involved in high affinity, proton-coupled transport of metabolites (particularly lactate) into neurons and may play a role in neuronal uptake of energy substrates released by glia (Itoh et al., 2003; Pellerin, 2003). MCT2 is localised to the post-synaptic compartment in many human neurons and may have a specialised role in synaptic functioning Chiry et al., 2008, Pierre et al., 2009,). Regulation of *SLC16A7* has also been linked to disorders of the brain: loss or under-expression has been associated with temporal-lobe epilepsy (Lauritzen et al., 2012) and it may be expressed and methylated at different levels in psychotic patients versus controls (Chen et al., 2014).

### MDD-Associated SNPs

The SNP on chromosome 1 associated with MDD only was located close to *LRRIQ3* and *FPGT*. *LRRIQ3* encodes leucine-rich repeat (LRR) and IQ motif containing protein 3, a calcium-channel component. LRR-domain containing proteins in general are involved in cell-cell communication, including processes involved in innate immunity and neuronal development (Bella, Hindle, McEwan, & Lovell, 2008; Ng et al., 2011). *FPGT* encodes fucose-1-phosphate guanylyltransferase, a protein involved in the alternative (salvage) pathways of fucose metabolism (Becker & Lowe, 2003). Fucose metabolism is important in a variety of cell-cell communication and host-microbe interaction situations, but is also important during neuronal development (Becker & Lowe, 2003). Variants in the *LRRIQ3* region have been previously associated with schizophrenia (Ripke et al., 2014), neurodevelopmental disorders (Reuter et al., 2017) and migraine (Gormley et al., 2016).

### Pleiotropic SNPs in *LRFN5*

The significantly associated SNPs on chromosome 14 found within or close to *LRFN5* were either associated with MDD only or were pleiotropic and associated with both CPG and MDD. Due to the LD-based pruning carried out before the cFDR analysis, the MDD-only and pleiotropic SNPs are not in strong linkage disequilibrium and may thus be tagging different functional variants, potentially acting on risk in different ways. *LRFN5* encodes leucine-rich repeat (LRR) and fibronectin type 3 domain-containing protein 5. Proteins in the LRFN family span the plasma membrane, and their extracellular domains are thought to participate in the cell-cell interactions necessary for both neuronal development (Morimura, Inoue, Katayama, & Aruga, 2006; Nam, Mah, & Kim, 2011) and synapse formation (Choi et al., 2016). Lrfn5, along with another member of the Lrfn protein family, Lrfn2, may induce both inhibitory and excitatory presynaptic differentiation in nearby.(Lin et al 2018), a process that may play a critical role in brain development and function (Cordova-Palomera et al 2016). As a family, these genes are primarily expressed in the central nervous system. Polymorphic markers within or close to *LRFN5* have been reported to be associated with progressive autism and familial schizophrenia (De Bruijn et al., 2010; B. Xu et al., 2009). Reduced expression of Lrfn5 has also been reported to contribute to neuro-inflammation (Zhu et al., 2016). Each of the four pleiotropic SNPs has opposing directions of effect in MDD and CPG. For example, the effect allele at rs11846556 is associated with an increase in CPG but with reduced risk of MDD. This may be explained by theories whereby structural and connectivity-related changes in the brain which facilitate development and maintenance of chronic pain involve neurogenesis and synaptic plasticity (Apkarian et al., 2011; Baliki, Mansour, Baria, & Apkarian, 2014; Vasic & Schmidt, 2017), but impaired neurogenesis is associated with depression (Fang, Demic, & Cheng, 2018; Jacobs, van Praag, & Gage, 2000). Additionally, differing expression patterns of *LRFN5* in the brain and CNS vs the periphery may be involved in opposing direction of effect associated with pleiotropic SNPs.

### Cis-eQTLs

SNPs within and close to *LRFN5* on chromosome 14, and in the region close to long non-coding RNA gene *LINC01360* on chromosome 1, were found to be cis-eQTLs. rs35641559 on chromosome 1, associated with MDD only, was found to be significantly associated with *LOC105378800* expression levels in the testis. All four pleiotropic SNPs on chromosome 14, were found to be significantly associated with *LRFN5* expression levels in a range of tissues, including the brain. Further investigation of potential regulatory roles for these or other nearby variants is warranted.

### Role of rs11846556 in the biology of chronic pain and depression

Number of A alleles at rs11846556, one of the pleiotropic SNPs identified here, was found to be associated with risk of chronic pain (and with higher values of the chronic pain category phenotype), but not with MDD risk or severity (in the UK Biobank cohort), and the effect size for the chronic pain association was not attenuated by inclusion of MDD in the regression model. This relationship is consistent with mediated pleiotropy rather than biological pleiotropy, i.e. a pathway in which genotype influences chronic pain risk, which in turn affects MDD risk, rather than influencing both by separate mechanisms. Formal mediation analyses was beyond the scope of this study primarily due to probable sequential ignorability violation and unmeasured confounders (Daniel, De Stavola, Cousens, & Vansteelandt, 2015; Imai, Keele, & Tingley, 2010).

As discussed above, each of the 4 SNPs tagged as pleiotropic, including rs11846556, are also cis-eQTLs for *LRFN5*. The SNP we explored in more detail, rs11846556, is a cis-eQTL of *LRFN5* in the cerebellum and cerebral hemisphere, along with tibial artery (GTEx single-tissue eQTL lookup). The number of A alleles at this SNP is significantly associated with increased expression of *LRFN5* in CNS tissues, but with a trend towards decreased expression in peripheral tissues (Supplementary Figures 1a-d), suggesting the existence of tissue-specific regulatory effects. This SNP is also a cis-eQTL for expression of a long intergenic non-coding RNA (Gencode ID: ENSG00000258636.1) on the reverse strand with respect to *LRFN5,* but only in spleen and aorta.

Alterations in synaptic plasticity and neurogenesis in the hippocampus, which is often non-normal in cases of neuropathic pain, may underlie some of the cognitive and affective deficits seen in patients with major depression (Fasick et al 2015). However, in cases where depression is a downstream consequence of chronic pain, its onset may be influenced by lifestyle and other ‘above-the-skin’ factors that also operate downstream of chronic pain, rather than as a more direct result of synaptic changes related to *LRFN5* expression levels.

Our data therefore suggest that rs11846556 genotype is significantly associated with chronic pain, but not with MDD, and that the relationship between chronic pain and genotype is not attenuated by MDD - pleiotropic effects of rs11846556 (or the functional variant it is tagging) on chronic pain and MDD appear to be mediated rather than biological. Possession of more copies of the A allele at this locus is associated with increased *LRFN5* expression levels in the CNS, which may exert its effect on transition to a chronic pain state after a prior insult through consequent alterations to brain network connectivity.

### Strengths and Limitations

A key strength of this study lies in its enhanced power, through the use of the cFDR approach, to detect the contribution of genetic variants to chronic pain relative to previous genome-wide approaches. Limitations include the self-reported nature of the chronic pain phenotype and the somewhat general phrasing of the questions used to elicit data about this phenotype in the 23&Me and UK Biobank studies, which precluded a finer dissection of the nature and severity of the pain being reported. In addition, the questions asked in the two studies were slightly different, generating chronic pain measures that may be capturing somewhat different aspects of the overall phenotype. ‘Chronic pain grade’ as assessed in the Pfizer-23andMe GWAS may be capturing a more direct severity-related phenotype, whereas ‘chronic pain category’ as assessed in UKB is likely to be capturing a different aspect of chronic pain susceptibility related to sensitivity of the response to multiple insults, or to propensity to report. Despite this subtle distinction, we were able to demonstrate that an *LRFN5* SNP is associated with both these phenotypic pain measures. Future studies building on these findings would benefit from using clinically validated or supported phenotyping data, however.

## Conclusions

To date no common variants have been associated with presence of chronic pain at any site in the body or with number of sites affected or pain severity. In this study six novel SNPs were found to be associated with chronic pain grade, a measure of pain severity. cFDR analyses also increased power to find MDD-associated variants in comparison to genome-wide significance thresholds. Conjunctional cFDR (ccFDR) analyses revealed evidence of pleiotropic effects of variants in *LRFN5*. The regions around the significant SNPs contain genes involved in, amongst other processes, neuronal cell metabolism and development, innate immunity and cell-cell interaction. These regions have also been previously implicated in a range of neurodevelopmental and psychiatric disorders. One of the pleiotropic SNPs identified on chromosome 14 (rs11846556), which is a cis-eQTL of *LRFN5* in cerebellum and cerebral hemisphere, was found to be associated with chronic pain but not MDD in UK Biobank. This provides intriguing preliminary evidence that the pleiotropy associated with variation in *LRFN5* may be mediated rather than horizontal.

## Acknowledgements

We would like to thank the research participants and employees of 23andMe for making this work possible. We also thank all participants in the UK Biobank study. UK Biobank was established by the Wellcome Trust, Medical Research Council, Department of Health, Scottish Government and Northwest Regional Development Agency. UK Biobank has also had funding from the Welsh Assembly Government and the British Heart Foundation. Data collection was funded by UK Biobank.

RJS is supported by a UKRI Innovation-HDR-UK Fellowship (MR/S003061/1). JW is supported by the JMAS Sim Fellowship for depression research from the Royal College of Physicians of Edinburgh (173558). KJAJ is supported by an MRC Doctoral Training Programme Studentship at the Universities of Glasgow and Edinburgh. DJS acknowledges the support of the Brain and Behavior Research Foundation (Independent Investigator Award 1930), a Lister Prize Fellowship (173096) and the MRC Mental Health Data Pathfinder Award (MC_PC_17217).

## References

Aguet, F. et al. (2017) ‘Genetic effects on gene expression across human tissues’, Nature, 550(7675), pp. 204–213. doi: 10.1038/nature24277. URL: https://www.gtexportal.org/home/browseEqtls?location=chr1:1-50,000,000.

Andreassen, O. A. et al. (2013) ‘Improved Detection of Common Variants Associated with Schizophrenia and Bipolar Disorder Using Pleiotropy-Informed Conditional False Discovery Rate’, PLoS Genetics, 9(4). doi: 10.1371/journal.pgen.1003455.

Andreassen, O. A. et al. (2013) ‘Improved detection of common variants associated with schizophrenia by leveraging pleiotropy with cardiovascular-disease risk factors’, American Journal of Human Genetics. The American Society of Human Genetics, 92(2), pp. 197–209. doi: 10.1016/j.ajhg.2013.01.001.

Andreassen, O. A. et al. (2014) ‘Identifying common genetic variants in blood pressure due to polygenic pleiotropy with associated phenotypes’, Hypertension, 63(4), pp. 819–826. doi: 10.1161/HYPERTENSIONAHA.113.02077.

Apkarian, A. V., Hashmi, J. A., & Baliki, M. N. (2011). Pain and the brain: Specificity and plasticity of the brain in clinical chronic pain. Pain, 152(SUPPL.3), S49–S64. https://doi.org/10.1016/j.pain.2010.11.010

Benjamini, Y. and Hochberg, Y. (1995) ‘Controlling the false discovery rate: a practical and powerful approach to multiple testing’, Journal of the Royal Statistical Society, pp. 289–300. doi: 10.2307/2346101.

Baliki, M. N. et al. (2012) ‘Corticostriatal functional connectivity predicts transition to chronic back pain’, Nature Neuroscience. Nature Publishing Group, 15(8), pp. 1117–1119. doi: 10.1038/nn.3153.

Baliki, M. N. et al. (2014) ‘Functional reorganization of the default mode network across chronic pain conditions’, PLoS ONE, 9(9). doi: 10.1371/journal.pone.0106133.

Baliki, M. N. and Apkarian, A. V. (2015) ‘Nociception, Pain, Negative Moods, and Behavior Selection’, Neuron. Elsevier Inc., 87(3), pp. 474–491. doi: 10.1016/j.neuron.2015.06.005.

Becker, D. J. and Lowe, J. B. (2003) ‘Fucose: Biosynthesis and biological function in mammals’, Glycobiology, 13(7). doi: 10.1093/glycob/cwg054.

Bella, J. et al. (2008) ‘The leucine-rich repeat structure’, Cellular and Molecular Life Sciences, 65(15), pp. 2307–2333. doi: 10.1007/s00018-008-8019-0.

Chamberlain, Ushey and Zhu (2016). rsnps: Get ‘SNP’ (‘Single-Nucleotide’ ‘Polymorphism’) Data on the Web. R package version 0.2.0. https://CRAN.R-project.org/package=rsnps

Chen, Y., Hor, H. H., & Tang, B. L. (2012). AMIGO is expressed in multiple brain cell types and may regulate dendritic growth and neuronal survival. Journal of Cellular Physiology, 227(5), 2217–2229. https://doi.org/10.1002/jcp.22958

Chen, C. et al. (2014) ‘Correlation between DNA methylation and gene expression in the brains of patients with bipolar disorder and schizophrenia’, Bipolar Disorders, 16(8), pp. 790–799. doi: 10.1111/bdi.12255.

Choi, Y. et al. (2016) ‘SALM5 trans-synaptically interacts with LAR-RPTPs in a splicing-dependent manner to regulate synapse development’, Scientific Reports. Nature Publishing Group, 6(December 2015), pp. 1–12. doi: 10.1038/srep26676.

Córdova-Palomera, A., Tornador, C., Falcón, C., Bargalló, N., Brambilla, P., Crespo-Facorro, B., … Faanás, L. (2016). Environmental factors linked to depression vulnerability are associated with altered cerebellar resting-state synchronization. Scientific Reports, 6(March), 1–11. https://doi.org/10.1038/srep37384

Daniel, R. M. et al. (2015) ‘Causal mediation analysis with multiple mediators’, Biometrics, 71(1), pp. 1–14. doi: 10.1111/biom.12248.

Davis, K. A. S., Coleman, J. R. I., Adams, M., Allen, N., Breen, G., Cullen, B., … Hotopf, M. (2018). Mental health in UK Biobank: development, implementation and results from an online questionnaire completed by 157 366 participants. BJPsych Open, 4(03), 83–90. https://doi.org/10.1192/bjo.2018.12

Debernardi, R. et al. (2003) ‘Cell-specific expression pattern of monocarboxylate transporters in astrocytes and neurons observed in different mouse brain cortical cell cultures’, J Neurosci Res, 73(2), pp. 141–155. doi: 10.1002/jnr.10660.

Efron, B. (2007) ‘Size, power and false discovery rates’, Annals of Statistics, 35(4), pp. 1351–1377. doi: 10.1214/009053606000001460.

Fang, J., Demic, S., & Cheng, S. (2018). The reduction of adult neurogenesis in depression impairs the retrieval of new as well as remote episodic memory. PLoS ONE, 13(6), 1–23. https://doi.org/10.1371/journal.pone.0198406

Fasick, V., Spengler, R. N., Samankan, S., Nader, N. D., & Ignatowski, T. A. (2015). The hippocampus and TNF: Common links between chronic pain and depression. Neuroscience and Biobehavioral Reviews. https://doi.org/10.1016/j.neubiorev.2015.03.014

Flint, J. and Kendler, K. S. (2014) ‘The Genetics of Major Depression’, Neuron. doi: 10.1016/j.neuron.2014.01.027.

Gandy, K., Kim, S., Sharp, C., Dindo, L., Maletic-Savatic, M., & Calarge, C. (2017). Pattern separation: A potential marker of impaired hippocampal adult neurogenesis in major depressive disorder. Frontiers in Neuroscience, 11(OCT). https://doi.org/10.3389/fnins.2017.00571

Goto-Ito, S., Yamagata, A., Sato, Y., Uemura, T., Shiroshima, T., Maeda, A., … Fukai, S. (2018). Structural basis of trans-synaptic interactions between PTPδ and SALMs for inducing synapse formation. Nature Communications, 9(1), 1–9. https://doi.org/10.1038/s41467-017-02417-z

Gormley, P. et al. (2016) ‘Susceptibility Loci for Migraine’, Nature Genetics, 48(8), pp. 16–18. doi: 10.1038/ng.3598.

Gratten, J. and Visscher, P. M. (2016) ‘Genetic pleiotropy in complex traits and diseases: implications for genomic medicine’, Genome Medicine. Genome Medicine, 8(1), p. 78. doi: 10.1186/s13073-016-0332-x.

Greene, S. A. (2010) ‘Chronic Pain: Pathophysiology and Treatment Implications’, Topics in Companion Animal Medicine. Elsevier Inc., 25(1), pp. 5–9. doi: 10.1053/j.tcam.2009.10.009.

Halestrap, A. P. (2012) ‘The monocarboxylate transporter family-Structure and functional characterization’, IUBMB Life, 64(1), pp. 1–9. doi: 10.1002/iub.573.

Hashmi, J. A. et al. (2013) ‘Shape shifting pain: Chronification of back pain shifts brain representation from nociceptive to emotional circuits’, Brain, 136(9), pp. 2751–2768. doi: 10.1093/brain/awt211.

Hocking, L. J. et al. (2010) ‘Genetic variation in the beta2-adrenergic receptor but not catecholamine-O-methyltransferase predisposes to chronic pain: Results from the 1958 British Birth Cohort Study’, Pain. International Association for the Study of Pain, 149(1), pp. 143–151. doi: 10.1016/j.pain.2010.01.023.

Imai, K., Keele, L. and Tingley, D. (2010) ‘A General Approach to Causal Mediation Analysis’, Psychological Methods, 15(4), pp. 309–334. doi: 10.1037/a0020761.

Itoh, Y. et al. (2003) ‘Dichloroacetate effects on glucose and lactate oxidation by neurons and astroglia in vitro and on glucose utilization by brain in vivo’, Proceedings of the National Academy of Sciences, 100(8), pp. 4879–4884. doi: 10.1073/pnas.0831078100.

Jacobs, B., van Praag, H., & Gage, F. (2000). Adult brain neurogenesis and psychiatry a novel theory of depression. Mol Psychiatry, May(5(3)), 262–269.

Kawato, M., Kuroda, S., & Schweighofer, N. (2011). Cerebellar supervised learning revisited: Biophysical modeling and degrees-of-freedom control. Current Opinion in Neurobiology, 21(5), 791–800. https://doi.org/10.1016/j.conb.2011.05.014

Kent WJ, Sugnet CW, Furey TS, Roskin KM, Pringle TH, Zahler AM, Haussler D. The human genome browser at UCSC. Genome Res. 2002 Jun;12(6):996–1006.

Kent WJ, Hsu F, Karolchik D, Kuhn RM, Clawson H, Trumbower H, Haussler D. Exploring relationships and mining data with the UCSC Gene Sorter. Genome Res. 2005 May;15(5):737–41.

Lauritzen, F. et al. (2012) ‘Redistribution of monocarboxylate transporter 2 on the surface of astrocytes in the human epileptogenic hippocampus’, Glia, 60(7), pp. 1172–1181. doi: 10.3174/ajnr.A1256.Functional.

Lee, S., Lee, J. and Lee, D. (2014) ‘The genetic variation in Monocarboxylic acid transporter 2 (MCT2) has functional and clinical relevance with male infertility’, Asian Journal of Andrology, 16(5), pp. 694. doi: 10.4103/1008-682X.124561.

Liley, J. and Wallace, C. (2015) ‘A pleiotropy-informed Bayesian false discovery rate adapted to a shared control design finds new disease associations from GWAS summary statistics’, PLoS genetics, 11(2), p. e1004926. doi: 10.1371/journal.pgen.1004926.

Lin, Z., Liu, J., Ding, H., Xu, F., & Liu, H. (2018). Structural basis of SALM5-induced PTPδ dimerization for synaptic differentiation. Nature Communications, 9(1). https://doi.org/10.1038/s41467-017-02414-2

Macfarlane, G. J., Barnish, M. S. and Jones, G. T. (2017) ‘Persons with chronic widespread pain experience excess mortality: longitudinal results from UK Biobank and meta-analysis’, Annals of the Rheumatic Diseases, 76(11), pp. 1815–1822. doi: 10.1136/annrheumdis-2017-211476.

Mansour, A. R., Baliki, M. N., Huang, L., Torbey, S., Herrmann, K. M., Schnitzer, T. J., & Vania Apkarian, A. (2013). Brain white matter structural properties predict transition to chronic pain. Pain, 154(10), 2160–2168. https://doi.org/10.1016/j.pain.2013.06.044

McIntosh, A. M. et al. (2016) ‘Genetic and Environmental Risk for Chronic Pain and the Contribution of Risk Variants for Major Depressive Disorder: A Family-Based Mixed-Model Analysis’, PLoS Medicine, 13(8), pp. 1–17. doi: 10.1371/journal.pmed.1002090.

Merskey, H. and Bogduk, N. (1994) Classification of Chronic Pain, IASP Pain Terminology. doi: 10.1002/ana.20394.

Mogil, J. S. (2012) ‘Pain genetics: Past, present and future’, Trends in Genetics. Elsevier Ltd, 28(6), pp. 258–266. doi: 10.1016/j.tig.2012.02.004.

Morimura, N. et al. (2006) ‘Comparative analysis of structure, expression and PSD95-binding capacity of Lrfn, a novel family of neuronal transmembrane proteins’, Gene, 380(2), pp. 72–83. doi: 10.1016/j.gene.2006.05.014.

Nam, J., Mah, W. and Kim, E. (2011) ‘The SALM/Lrfn family of leucine-rich repeat-containing cell adhesion molecules’, Seminars in Cell and Developmental Biology. Elsevier Ltd, 22(5), pp. 492–498. doi: 10.1016/j.semcdb.2011.06.005.

Ng, A. C. Y. et al. (2011) ‘Human leucine-rich repeat proteins: a genome-wide bioinformatic categorization and functional analysis in innate immunity’, Proceedings of the National Academy of Sciences, 108(Supplement_1), pp. 4631–4638. doi: 10.1073/pnas.1000093107.

Pellerin, L. (2003) ‘Lactate as a pivotal element in neuron-glia metabolic cooperation’, Neurochemistry International, 43(4–5), pp. 331–338. doi: 10.1016/S0197-0186(03)00020-2.

Perneger Thomas V. ‘What’s wrong with Bonferroni adjustments’ BMJ 1998; 316 :1236

Pértega-Gomes, N. et al. (2013) ‘Monocarboxylate transporter 2 (MCT2) as putative biomarker in prostate cancer’, Prostate, 73(7), pp. 763–769. doi: 10.1002/pros.22620.

Pierre, K., Magistretti, P. J. and Pellerin, L. (2002) ‘MCT2 is a major neuronal monocarboxylate transporter in the adult mouse brain’, J Cereb Blood Flow Metab, 22(5), pp. 586–595. doi: 10.1097/00004647-200205000-00010.

Raymond, J. L., & Medina, J. F. (2018). Computational Principles of Supervised Learning in the Cerebellum. Annual Review of Neuroscience, 41(1), 233–253. https://doi.org/10.1146/annurev-neuro-080317-061948

Reuter, M. S. et al. (2017) ‘Diagnostic yield and novel candidate genes by exome sequencing in 152 consanguineous families with neurodevelopmental disorders’, JAMA Psychiatry, 74(3), pp. 293–299. doi: 10.1001/jamapsychiatry.2016.3798.

Ripke et al (2014) ‘Biological Insights From 108 Schizophrenia-Associated Genetic Loci’, Nature, 511(7510), pp. 421–427. doi: 10.1038/nature13595.Biological.

Rothman, K. J. (1990) ‘No Adjustments Are Needed for Multiple Comparisons’, Epidemiology, 1(1), pp. 43–46.

Sivakumaran, S. et al. (2011) ‘Abundant pleiotropy in human complex diseases and traits’, American Journal of Human Genetics. The American Society of Human Genetics, 89(5), pp. 607–618. doi: 10.1016/j.ajhg.2011.10.004.

Storey, J. D. (2002) ‘A direct approach to false discovery rates’, Journal of the Royal Statistical Society. Series B: Statistical Methodology, 64(3), pp. 479–498. doi: 10.1111/1467-9868.00346.

Suri, P. et al. (2018) ‘Genome-wide meta-analysis of 158,000 individuals of European ancestry identifies three loci associated with chronic back pain’, 48(7), pp. 1–23. doi: 10.1038/ng.3582.

Valença, I. et al. (2015) ‘Localization of MCT2 at peroxisomes is associated with malignant transformation in prostate cancer’, Journal of Cellular and Molecular Medicine, 19(4), pp. 723–733. doi: 10.1111/jcmm.12481.

Vasic, V. and Schmidt, M. H. H. (2017) ‘Resilience and vulnerability to pain and inflammation in the hippocampus’, International Journal of Molecular Sciences, 18(4). doi: 10.3390/ijms18040739.

Von Korff, M. et al. (1992) ‘Grading the severity of chronic pain.’, Pain, 50(1092), pp. 133–49. doi: 10.1016/0304-3959(92)90154-4.

Vos, T. et al. (2015) ‘Global, regional, and national incidence, prevalence, and years lived with disability for 301 acute and chronic diseases and injuries in 188 countries, 1990-2013: A systematic analysis for the Global Burden of Disease Study 2013’, The Lancet. doi: 10.1016/S0140-6736(15)60692-4.

Wang, X. F. et al. (2017) ‘Linking Alzheimer’s disease and type 2 diabetes: Novel shared susceptibility genes detected by cFDR approach’, Journal of the Neurological Sciences. Elsevier B.V., 380, pp. 262–272. doi: 10.1016/j.jns.2017.07.044.

Wray, N. R. et al. (2018) ‘Genome-wide association analyses identify 44 risk variants and refine the genetic architecture of major depression.’, Nature genetics, 50(May), p. 167577. doi: 10.1038/s41588-018-0090-3.

Xu, Q., & Yaksh, T. L. (2011). A brief comparison of the pathophysiology of inflammatory versus neuropathic pain. Current Opinion in Anaesthesiology, 24(4), 400–407. https://doi.org/10.1097/ACO.0b013e32834871df

Yu, W., & Krook-Magnuson, E. (2015). Cognitive Collaborations: Bidirectional Functional Connectivity Between the Cerebellum and the Hippocampus. Frontiers in Systems Neuroscience, 9(December), 1–10. https://doi.org/10.3389/fnsys.2015.00177

Zhu, Y., Yao, S., Augustine, M. M., Xu, H., Wang, J., Sun, J., … Chen, L. (2016). Neuron-specific SALM5 limits inflammation in the CNS via its interaction with HVEM. Science Advances, 2(4), e1500637. https://doi.org/10.1126/sciadv.1500637

Zorina-Lichtenwalter, K. et al. (2016) ‘Genetic predictors of human chronic pain conditions’, Neuroscience. The Authors, 338, pp. 36–62. doi: 10.1016/j.neuroscience.2016.04.041.

