## Supplemental Fig 1 for "Identification of novel common variants associated with chronic pain using conditional false discovery rate analysis with major depressive disorder and assessment of pleiotropic effects of *LRFN5*"

Figure 1: Boxplots from single-tissue eQTL lookups of rs11846556

a)

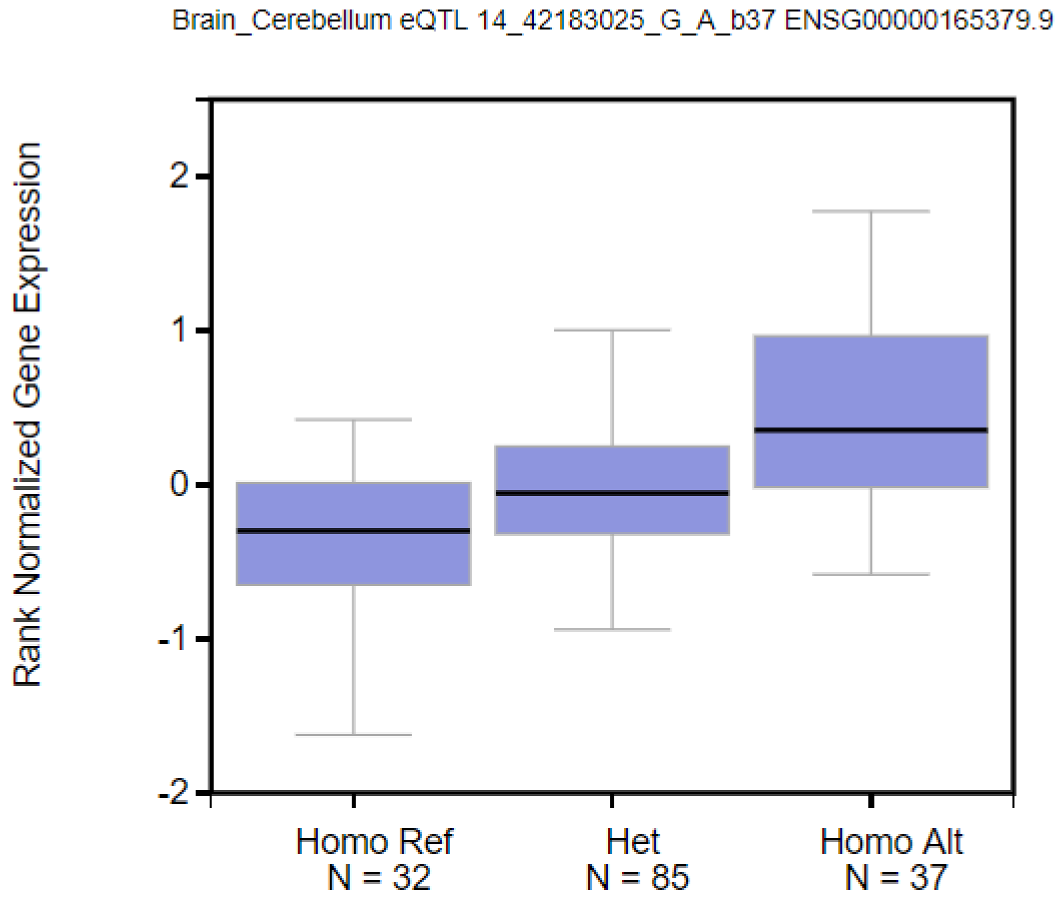

b)

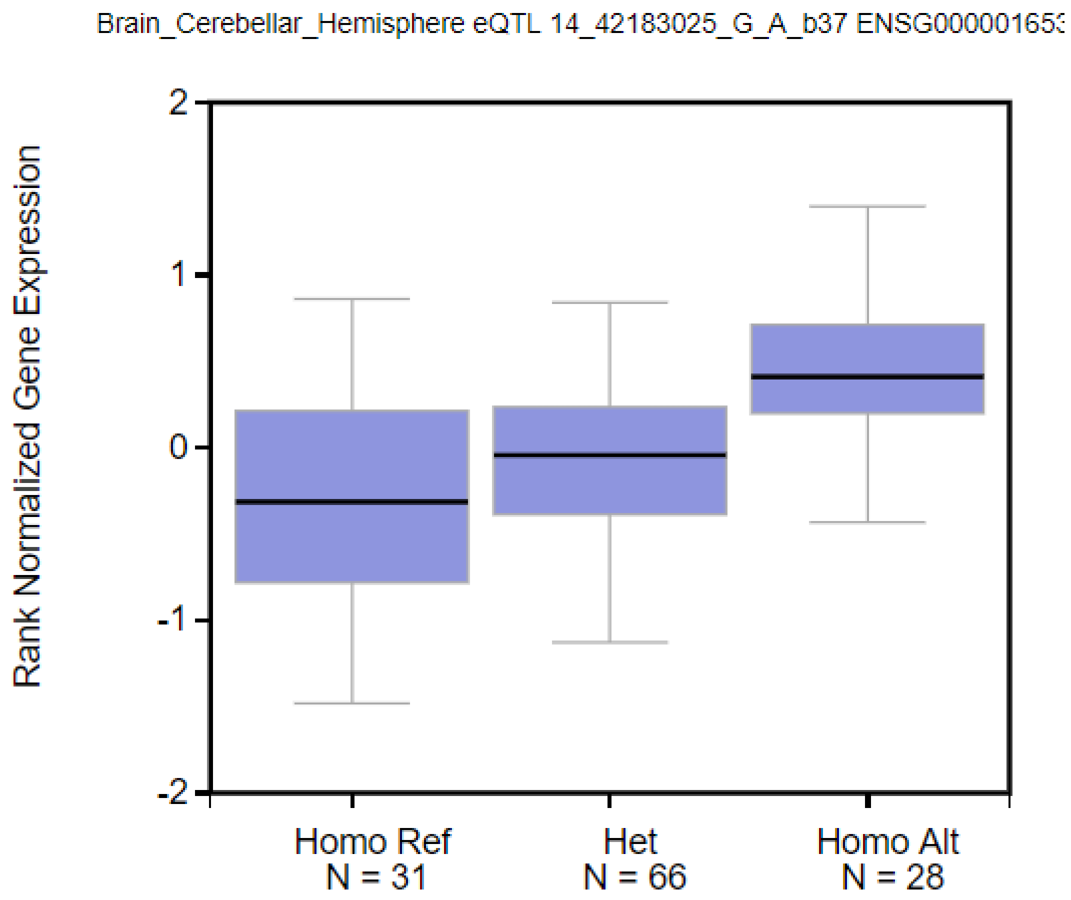
Brain_Cerebellar_Hemisphere eQTL 14_42183025_G_A_b37 ENSG00000165
Rank Normalized Gene Expression
2
1
0
-1
-2
Homo Ref
N = 31
Het
N = 66
Homo Alt
N = 28

c)

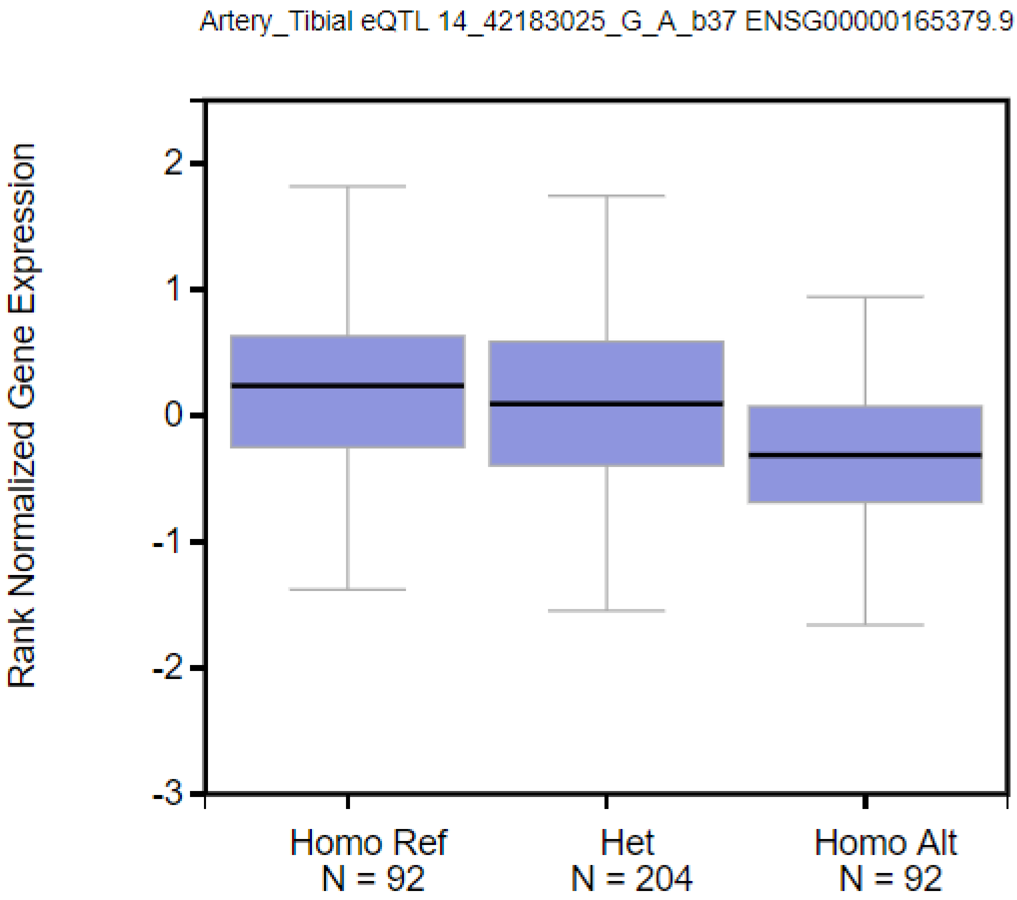

Artery_Tibial eQTL 14_42183025_G_A_b37 ENSG00000165379.9
Rank Normalized Gene Expression
2
1
0
-1
-2
-3
Homo Ref
N = 92
Het
N = 204
Homo Alt
N = 92

d)

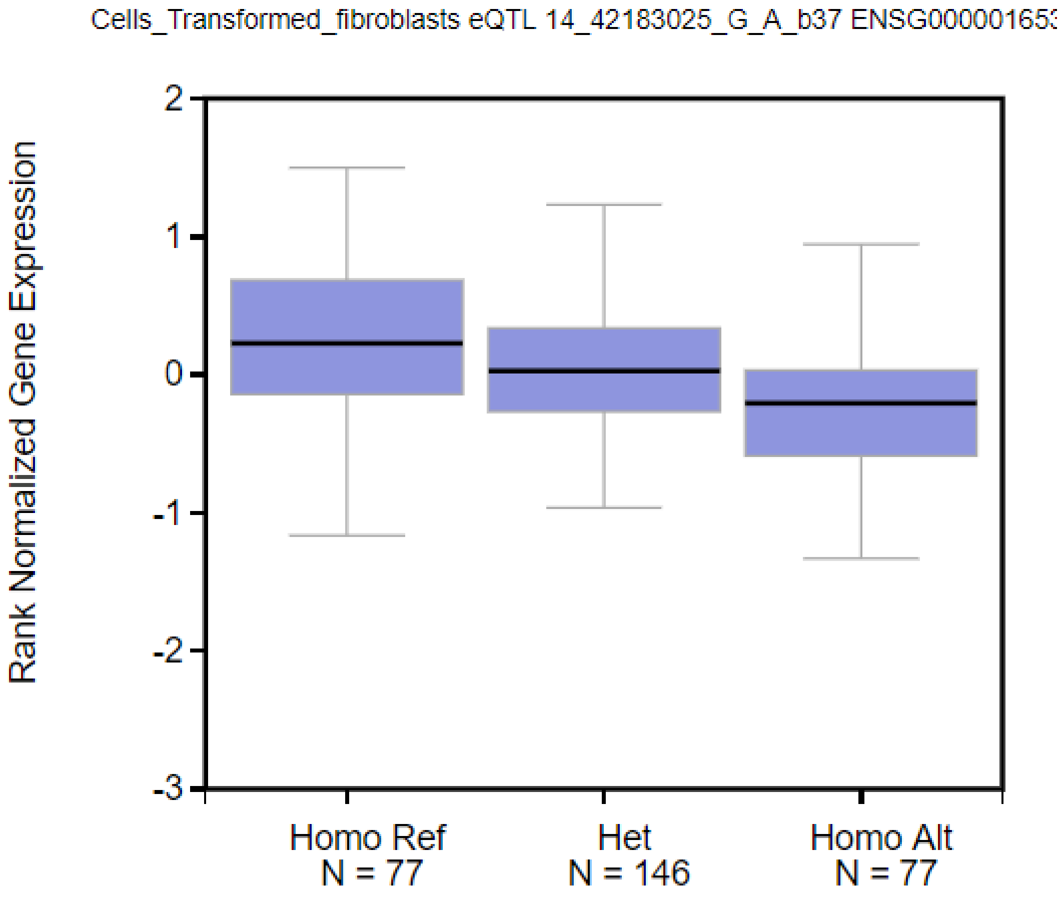


Figure 1: Boxplot output from single-tissue eQTL lookups (GTeX IGV eQTL Browser) of rs11846556, showing rank normalised gene expression values for *LRFN5.* A) Cerebellum B) Cerebellar hemisphere C) Tibial artery D) Transformed fibroblasts.
