## Supplemental Table 2c for "Identification of novel common variants associated with chronic pain using conditional false discovery rate analysis with major depressive disorder and assessment of pleiotropic effects of *LRFN5*"

**Table 2c: IGV eQTL Browser results.**

| rsID | Chrom | **cFDR-Associated Trait** | cis-eQTL Tissue Location(s) |
|---|---|---|---|
| rs35641559 | 1 | MDD | testis |
| rs149981001 | 12 | CPG | NA |
| rs147573737 | 12 | CPG | NA |
| rs4904790 | 14 | MDD | cerebellar hemisphere, cerebellum, transformed fibroblasts |
| rs1584317 | 14 | MDD | transformed fibroblasts |
| rs11846556 | 14 | Both | aorta, tibial artery, cerebellar hemisphere, cerebellum, transformed fibroblasts, spleen |
| rs10131184 | 14 | Both | subcutaneous adipose, aorta, tibial artery, cerebellar hemisphere, cerebellum, thyroid, transformed fibroblasts, oesophagus muscularis, ovary, skin (lower leg, not sun-exposed), spleen |
| rs8015100 | 14 | Both | omentum, aorta, coronary artery, cerebellum, cerebellar hemisphere, transformed fibroblasts, oesophagus muscularis, ovary, spleen, thyroid |
| rs11157241 | 14 | Both | subcutaneous adipose, aorta, tibial artery, cerebellar hemisphere, cerebellum, transformed fibroblasts, oesophagus muscularis, skin (lower leg, not sun-exposed), spleen, thyroid |

| | | | |
|---|---|---|---|
| rs10138559 | 14 | MDD | coronary artery, aorta, tibial artery, cerebellum, cerebellar hemisphere, transformed fibroblasts, oesophagus muscularis, spleen, thyroid |
| rs10872954 | 14 | MDD | aorta, transformed fibroblasts, spleen |

The tissue location(s) of cis-eQTLs where a gene is significantly regulated by the queried SNP (rsID column) (FDR < 0.05) are listed, along with SNP ID (rsID), chromosomal location (Chrom) and cFDR-associated trait.
