## Supplemental Table 2b for "Identification of novel common variants associated with chronic pain using conditional false discovery rate analysis with major depressive disorder and assessment of pleiotropic effects of *LRFN5*"

**Table 2b: UCSC Genome Browser Search Results**.

| rsID | Chromosome | cFDR-Associated Trait | Gene(s) |
|---|---|---|---|
| rs35641559 | 1 | MDD | *LINC01360, LRRIQ3, FPGT, FPGT-TNNI3K* |
| rs149981001 | 12 | CPG | *SLC16A7* |
| rs147573737 | 12 | CPG | *SLC16A7* |
| rs4904790 | 14 | MDD | *LRFN5* |
| rs1584317 | 14 | MDD | *LRFN5* |
| rs11846556 | 14 | Both | *LRFN5* |
| rs10131184 | 14 | Both | *LRFN5* |
| rs8015100 | 14 | Both | *LRFN5* |
| rs11157241 | 14 | Both | *LRFN5* |
| rs10138559 | 14 | MDD | *LRFN5* |
| rs10872954 | 14 | MDD | *LRFN5* |
