## Supplemental Table 2a for "Identification of novel common variants associated with chronic pain using conditional false discovery rate analysis with major depressive disorder and assessment of pleiotropic effects of *LRFN5*"

**Table 2a: Output of ‘rsnps’ query**

| rsID | Chromosome | cFDR-Associated Trait | Gene(s) | Alleles | Major | Minor | MAF | AA |
|---|---|---|---|---|---|---|---|---|
| rs35641559 | 1 | MDD | LOC105378800 | C/T | T | C | 0.4641 | T |
| rs149981001 | 12 | CPG | NA | C/T | C | T | 0.0022 | C |
| rs147573737 | 12 | CPG | NA | C/T | T | C | 0.0024 | T |
| rs4904790 | 14 | MDD | LRFN5 | C/T | C | T | 0.3175 | C |
| rs1584317 | 14 | MDD | LRFN5 | C/G | G | C | 0.2993 | G |
| rs11846556 | 14 | Both | LRFN5 | A/G | A | G | 0.3676 | G |
| rs10131184 | 14 | Both | LRFN5 | A/G | G | A | 0.239 | G |
| rs8015100 | 14 | Both | LRFN5 | A/T | A | T | 0.2546 | A |
| rs11157241 | 14 | Both | NA | C/T | T | C | 0.2508 | T |
| rs10138559 | 14 | MDD | NA | C/T | C | T | 0.4399 | T |
| rs10872954 | 14 | MDD | NA | A/G | A | G | 0.4343 | A |

SNP ID (rsID), location, cFDR-associated trait, associated genes (Gene(s)), minor allele frequency (MAF) and ancestral allele (AA) are shown. ‘NA’ indicates no result for that query in that category.
