## Supplemental Table 1 for "Identification of novel common variants associated with chronic pain using conditional false discovery rate analysis with major depressive disorder and assessment of pleiotropic effects of *LRFN5*"

**Table 1: Pfizer-23andMe Participant Demographic Information.**

| CPG level | Total | M | F | Age 0-30 | Age 30-45 | Age 45-60 | Age 60+ |
|---|---|---|---|---|---|---|---|
| 0 | 12758 | 7502 | 5256 | 3059 | 4347 | 2938 | 2414 |
| 1 | 5803 | 2809 | 2994 | 785 | 1435 | 1734 | 1849 |
| 2 | 2344 | 899 | 1445 | 253 | 520 | 782 | 789 |
| 3 | 1349 | 374 | 975 | 106 | 323 | 483 | 437 |
| 4 | 1047 | 302 | 745 | 57 | 231 | 421 | 338 |

N = 23,301 individuals were included in the Pfizer-23andMe chronic pain grade (CPG) GWAS. CPG level 0-4 = von Korff chronic pain grade 0-4, M = Male, F = Female.
